# Stress, novel sex genes and epigenetic reprogramming orchestrate socially-controlled sex change

**DOI:** 10.1101/481143

**Authors:** Erica V Todd, Oscar Ortega-Recalde, Hui Liu, Melissa S Lamm, Kim M Rutherford, Hugh Cross, Michael A Black, Olga Kardailsky, Jennifer A Marshall Graves, Timothy A Hore, John R Godwin, Neil J Gemmell

## Abstract

Bluehead wrasses undergo dramatic, socially-cued female to male sex change. We apply transcriptomic and methylome approaches in this wild coral reef fish to identify the primary trigger and subsequent molecular cascade of gonadal metamorphosis. Our data suggest that the environmental stimulus is exerted via the stress axis, that repression of the aromatase gene (encoding the enzyme converting androgens to estrogens) triggers a cascaded collapse of feminizing gene expression, and identifies notable sex-specific gene neofunctionalization. Furthermore, sex change involves distinct epigenetic reprogramming and an intermediate state with altered epigenetic machinery expression akin to the early developmental cells of mammals. These findings reveal at a molecular level how a normally committed developmental process remains plastic and is reversed to completely alter organ structures.

**One Sentence Summary:** Ovary to testis transformation in a sex-changing fish involves transcriptomic and epigenomic reprogramming.

## Main Text

In most organisms a fundamental dichotomy is established in early embryonic development; individuals become either female or male and maintain these fates throughout life. However, some plant and animal species exhibit remarkably diverse and plastic sexual developmental patterns (*1–3*), and some even retain the ability to change sex in adulthood (*4, 5).* Such functional sex change is widespread in marine fishes, appearing in 27 families (*6)*. Among the most outstanding, and well-studied, is the bluehead wrasse *(Thalassoma bifasciatum),* a small coral reef fish that undergoes rapid and complete female-to-male sex reversal in response to a social cue (*7*).

While the sex change process and its evolutionary advantages are now well known (*5, 8*), there remain longstanding questions about how environmental influences initiate such dramatic changes in sexual identity, and what molecular processes orchestrate this transformation (*9, 10*).

Across vertebrates, antagonism between core male- and female-promoting gene networks is now recognized as crucial to the establishment and maintenance of gonadal fate (*11*). Discoveries in mice that loss of FOXL2 transcription factor is sufficient to induce the transdifferentiation of mature ovary into testis (*12*), and that testicular cells will become ovarian cells if the transcription factor DMRT1 is lost (*13*), suggest gonadal bipotentiality is retained into adulthood (*14*), and presents a mechanism through which female–male and male–female gonadal sex reversal may be controlled in fish. A key target of both factors is the gene encoding aromatase (the enzyme responsible for estrogen production), the expression of which is shown to be environmentally sensitive (e.g., (*15*)) and when inhibited results in female-male sex reversal (e.g., (*16*)).

Epigenetic processes are also suspected to be key mediators and effectors of sex reversal (*15, 17*). Temperature-sensitive sex-reversal involves global methylation modification in the half-smooth tongue sole (18); with sex-specific methylation states of major sex-pathway genes inverted following sex reversal. Similarly, sex reversal in the dragon lizard (*19*), and temperature-dependent sex determination in red-eared slider turtles (*20*), appears to involve temperature-sensitive expression changes in epigenetic regulator genes of the Jumonji family, namely *Kdm6b* and *Jarid2.* Therefore, a change in methylation state may facilitate reprogramming of sexual fate at a cellular level (*11*).

Sex reversal in response to social cues, as seen in the bluehead wrasse, is an especially striking example of phenotypic and sexual plasticity (*4*). Bluehead wrasses begin their reproductive life as females, but routinely reverse sex in the absence of a socially dominant male (21). Removal of the dominant terminal-phase (TP) male from its territory induces rapid and complete sex change of the largest female (*7, 22*) (Fig. 1). This female exhibits behavioral changes within hours, and complete transformation of a mature ovary to functional testis in 8-10 days. The absence of differentiated male sexual tissue in the ovaries of bluehead wrasses implies that some form of cellular reprogramming underlies sex change in this species, either by a direct cellular transition (i.e., transdifferentiation) or via a de differentiated intermediate stage, rather than by alterations in the proportion of spermatogenic and oogenic cell populations as in the bisexual gonad or ovotestis of many other sex changing fishes (e.g., clownfish and bidirectional sex-changing gobies; reviewed by (*10*)).

**Fig. 1.**
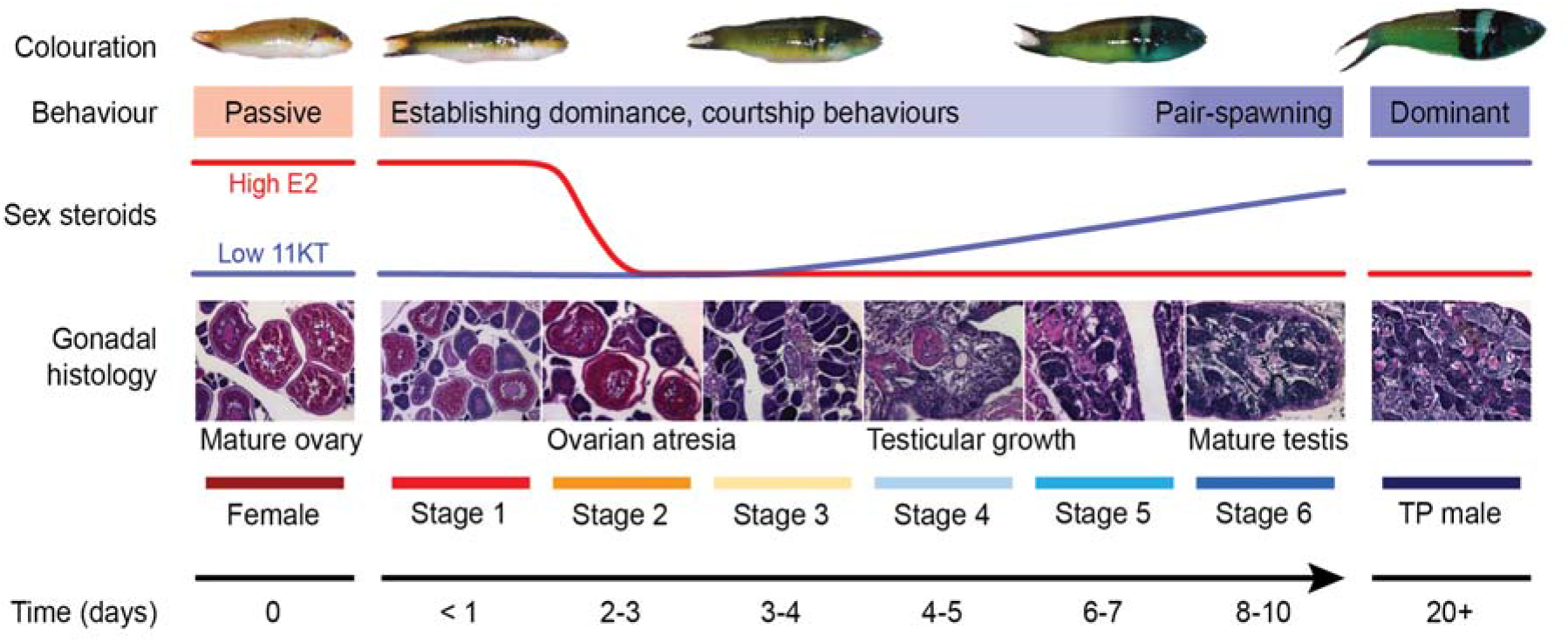
Sex change in the bluehead wrasse. Schematic of sex change summarizing changes in external coloration, behavior, serum steroid levels and gonadal histology across time. Within hours of removing TP males, the largest female displays aggression and male courtship behaviors, and adopts darker spawning coloration, but still possesses healthy ovaries (stage 1). Transitioning females establish dominance within 1-2 days, after which serum estrogen (E2) levels collapse and ovarian atresia is observable (stage 2). Ovarian atresia is advanced by 3-4 days (stage 3). Testicular tissues are observed by day 4-5 and serum 11-Ketotestosterone (11-KT) begins to rise (stage 4), before spermatogenesis begins by day 6-7 (stage 5). Within 8-10 days, transitioning fish are producing mature sperm and pair-spawning with females (stage 6). Full TP male coloration develops within ~20 days. Gonadal stages are classified according to (*23*). Hormonal changes are predicted based on patterns in the congener *Thalassoma duperrey (23*). For detailed descriptions of behavioral and morphological changes, see (*22*).

Here we applied transcriptomic and nucleotide-level methylation approaches to identify the primary trigger and subsequent molecular cascade that orchestrates gonad remodeling in the bluehead wrasse. Our results provide the first detailed molecular picture of female-to-male sex reversal in a vertebrate, and underscore the role of epigenetics and pluripotency in sex determination.

## Results

We induced wild female bluehead wrasses to change sex in the field by removing dominant TP males from established social groups on patch reefs off the Florida coast. We collected a time-series of brain and gonadal samples across the entire sex change process, assigning animals to 6 successive stages based on behaviors observed at time of capture (*22*) and gonadal histology (*23*) (Fig. 1). As controls we collected 6 small females that experienced the social manipulation but showed no signs of sex change (control females), and 8 dominant terminal phase (TP) males.

### Transcriptome-wide gene expression patterns across sex change

We quantified transcriptome-wide gene expression across sex change using RNA sequencing (RNA-seq), and the *de novo* assembled transcriptome from our previous study (*24*) as a reference. Expression patterns across samples were visualized using principal component analysis (PCA) (*25*). We observed very little variation in gene expression in the brain (Fig. 2A), consistent with other studies in teleosts showing limited sex bias in brain structure or gene expression (*26, 27*).

**Fig. 2.**
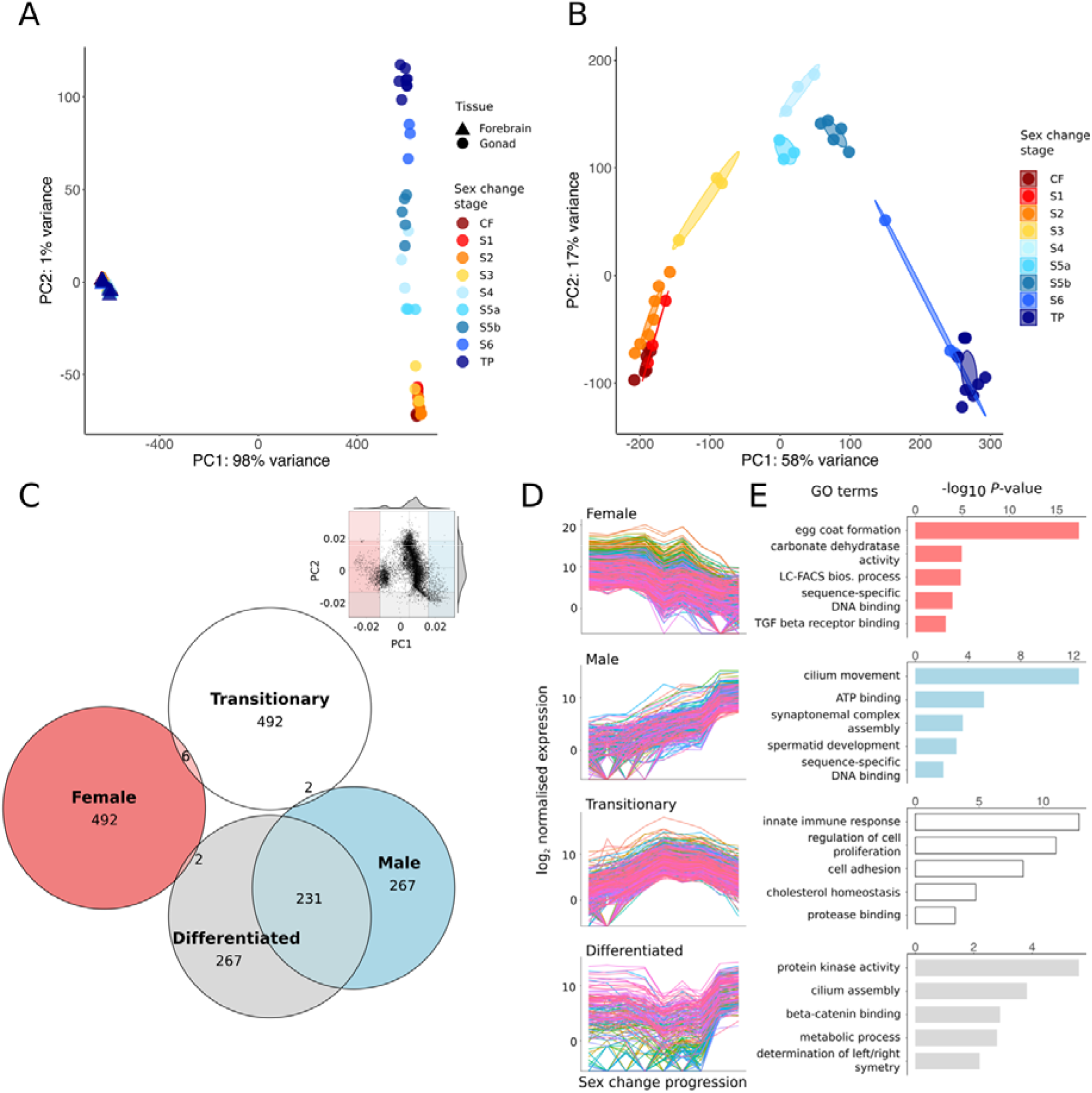
Global gene expression changes during sex change. **(A)** Principal component analysis (PCA) showing close clustering of brain samples but separation of gonad samples by sex change stage. **(B)** PCA (10,000 most variable genes) of gonad samples. The transition from ovary to testis is captured along PC1 (58% variance), whereas PC2 (17% variance) delineates fully differentiated gonads of control females (bottom left) and TP males (bottom right) from those of transitioning fish. **(C)** Inset, component loadings were used to identify transcripts contributing most to PC1 and PC2. Shaded sections define 5th and 95th percentiles defining four spatial regions: ‘Female’ (left), ‘Male’ (right), ‘Differentiated’ (bottom) and ‘Transitionary’ (top). The Euler diagram shows numbers of transcripts uniquely assigned to each region. **(D)** Expression patterns across sex change of transcripts uniquely assigned to each of the four spatial regions, showing four distinct patterns of expression: ‘Female’ (declining), ‘Male’ (increasing), ‘Transitionary’ (highest mid-sex change) and ‘Differentiated’ (lowest mid-sex change), and **(E)** representative GO terms for these transcripts. LC-FACS bios. process, long-chain fatty-acyl-CoA biosynthetic process; TGF beta receptor binding, Transforming growth factor beta receptor binding.

In contrast, striking variation in expression was observed in the gonad, and samples strongly clustered by sex change stage (Fig. 2B). Samples at stage 5 (ongoing spermatogenesis) formed two distinct clusters in the PCA and were subdivided into stage 5a and 5b in further analyses. Strikingly, gonad samples were clearly organized first by sexual development from female to male (horizontal axis), and second by developmental commitment (vertical axis) (Fig. 2B). This suggested that intermediate phases of sex change represent unique transitional cell types rather than simply different proportions of differentiated male and female tissues.

To investigate those transcripts contributing most to each extreme of PC1 (female, male) and PC2 (committed, transitionary), we used component loadings to select the top 500 transcripts driving each axis direction (Fig. 2C). This resulted in extensive overlap between the ‘male’ and ‘differentiated’ regions of the PCA. Visualizing expression of transcripts unique to each group (Fig. 2D) confirmed that the extremes of PC1 are characterized by transcripts that are female-biased and downregulated, or male-biased and upregulated across sex change, whereas PC2 is driven by transcripts with highest expression either in transitionary stages or sexually differentiated gonads. Gene Ontology (GO) terms (Fig. 2E) associate transcripts of the ‘female’ and ‘male’ regions with oocyte- and sperm-specific processes, respectively, and sex-specific transcription factor activity. Transcripts unique to the ‘transitionary’ region associated with innate immunity, protein catabolism, cell proliferation and adhesion processes, reflecting dynamic disassembly and rebuilding of the gonad at mid-sex change. The ‘differentiated’ region was associated with protein kinase activity, ß-catenin signaling, metabolic and developmental processes.

### Specific gene expression changes

Sex change involves transition from a female-biased expression landscape, to one that is male-biased. This transition is not linear, and is punctuated by large numbers of differentially expressed transcripts between stage 3 (advanced ovarian atresia) and 4 (early testicular development), and between stage 5 (early spermatogenesis) and 6 (spermiation) (Fig. S1). Classical ovary-specific genes *(nr0b1 (dax1), figla* and *gdf9*) became progressively downregulated as sex change progressed (Fig. 3).

**Fig. 3.**
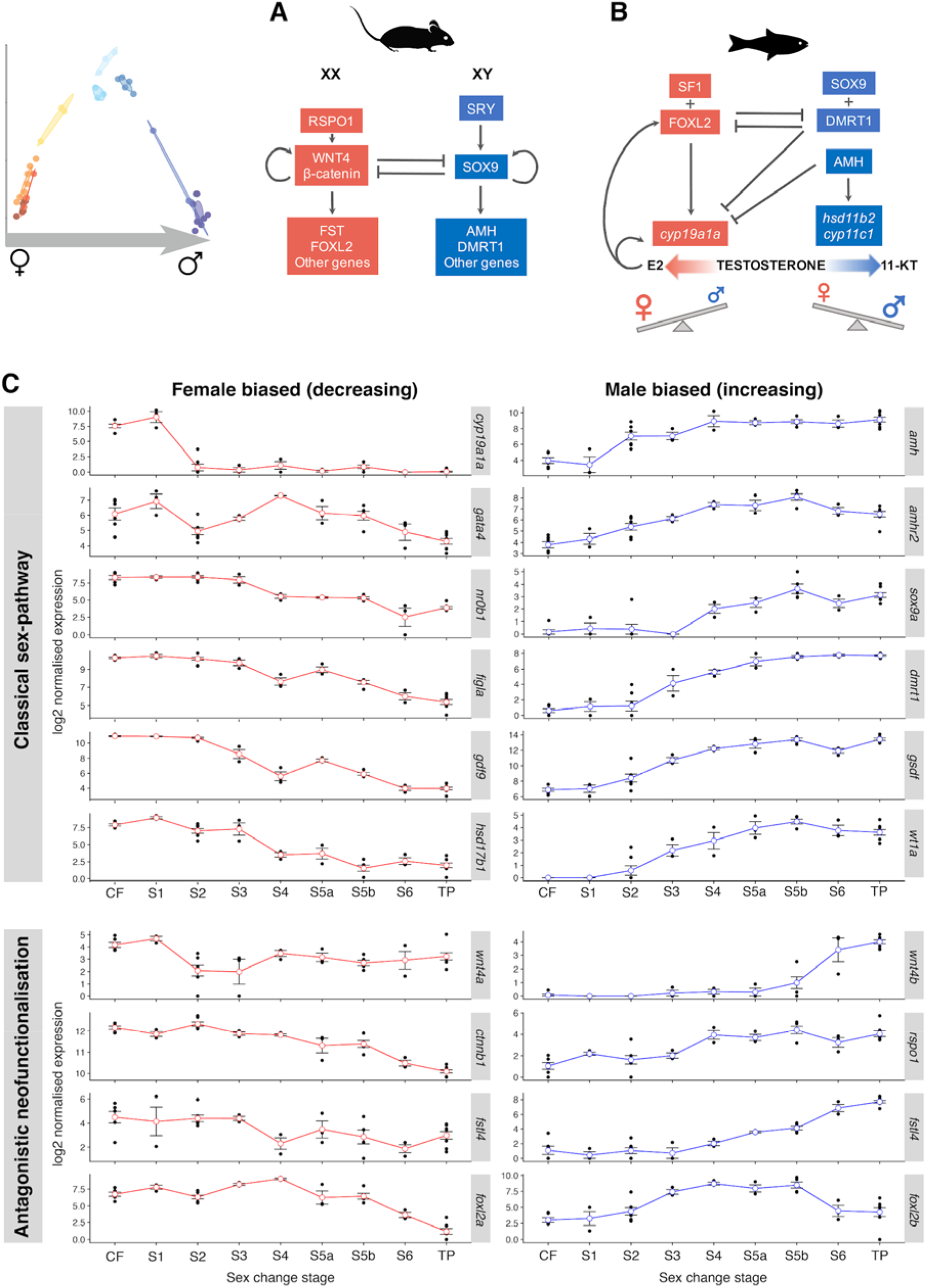
Sex change involves transition from female-to male-specific expression and gene neofunctionalization. **(A)** Mammalian model of sex determination and development. In males, SRY (sex-determining region Y) activates SOX9 to initiate male development, while blocking expression of feminization genes that would in turn antagonize masculinizing expression (*11*). **(B)** Teleost model of sexual development. Diverse factors determine sex in fishes, yet conserved downstream effectors act in feminizing and masculinizing pathways to promote female or male development, respectively, while antagonizing the opposing sexual pathway. Testosterone is a pro-hormone and is converted to estrogen (E2) by gonadal aromatase (encoded by *cyp19a1a)* to promote ovarian function, or to 11-Ketotestosterone (11-KT) by the products of *hsd11b2* and *cyp11c1* to promote testicular function. **(C)** Normalized expression of classical sex-pathway genes across sex change in bluehead wrasse gonads. At stage 2, *cyp19a1a* is sharply downregulated, after which feminizing expression collapses as female fish transition to males. Upregulation of masculinizing gene expression begins with *amh* and its receptor *amhr2.* Classically feminizing genes within the Respondin/Wnt/β-catenin signaling pathway *(wnt4a/b, rspo1,* and two transcripts annotated as *fstl4* are shown as examples), and also *foxl2,* are duplicated in the bluehead wrasse with orthologues showing testis-specific expression. CF, control female; S1-6, stage 1 to 6; TP, terminal phase male.

Testis-specific genes became activated from mid sex change. *Amh* (anti-Müllerian hormone) and its receptor *amh2* were the first male-pathway genes to show increased expression, at stage 2 and before the appearance of male tissues in the gonad, followed by other classical male-promoting genes (e.g., *dmrt1, sox9, gsdf)* (Fig. 3). Stages characterized by active testicular growth and spermatogenesis (stage 4 onwards) displayed increasing expression of *cyp11c1* and *hsd11b2,* consistent with their function to convert testosterone to the more potent teleost androgen 11-Ketotestosterone (11-KT) (Fig. 4).

**Fig. 4.**
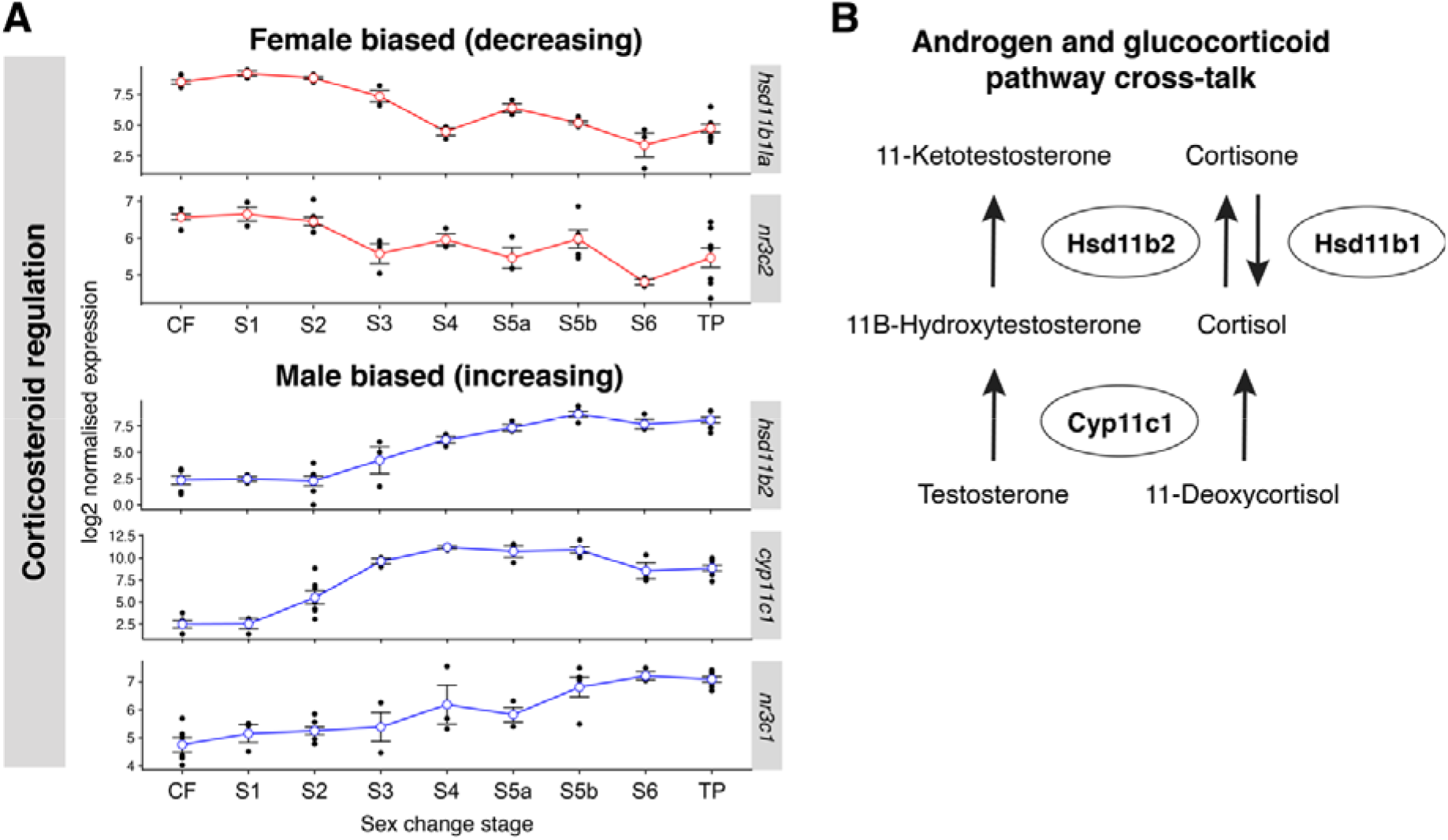
Dynamic sex-specific expression of androgenesis and glucocorticoid factors. **(A)** Normalized gonadal expression of androgenesis and cortisol pathway genes across sex change. Upregulation of *cyp11c1* at stage 2 implies increased local cortisol production. Then, changes in expression of 11β-Hsd enzymes at stage 3 suggest a shift from cortisone-cortisol regeneration *(hsd11b1a* downregulated) to cortisol-cortisone inactivation *(hsd11b2* upregulated). Concurrent upregulation of *hsd11b2* would also promote 11-Ketotestosterone (11-KT) synthesis, the most potent teleost androgen. Genes encoding the glucocorticoid *(nr3c1)* and mineralocorticoid *(nr3c2)* receptors show opposing sex-specific expression patterns that also imply highest cortisol activity at early sex change. These expression patterns suggest a window of high cortisol activity at stage 2 that is concurrent with arrested aromatase expression and ovarian atresia, followed by the stimulation of 11-KT production by stage 3. **(B)** Schematic showing cross-talk between the androgenesis and glucocorticoid pathways. The 11β-hydroxylase enzyme (Cyp11c1, homologous to mammalian CYP11B) and two 11β-hydroxysteroid dehydrogenases (Hsd11b1, Hsd11b2) are together responsible for 11-KT production, and the interconversion of cortisol. Cortisol is produced via the action of Cyp11c1, whereas 11β-Hsd enzymes control its interconversion with inactive cortisone, thus mediating the stress response. Cortisol itself stimulates 11β-Hsd expression, resulting in the production of both 11-KT and inactive cortisone.

To identify the most upstream effectors of sex change, we focused on expression changes in the earliest phases. Strikingly, *cyp19a1a* and *cyp19a1b* showed dramatic downregulation at early sex-change, in gonad and brain respectively, and remained at low levels thereafter. These genes encode the brain- and gonad-specific forms of aromatase, which converts testosterone to estradiol. The balance of estrogen (17*β*-estradiol, E2) versus androgen (11-KT) production is known to control sexual fate in teleost fish (*1, 28*) and aromatase inhibitors can effectively induce female-to-male sex reversal (*16, 29*). Our data provide the first whole-transcriptome evidence that aromatase downregulation is an early switch initiating sex change in both the brain and gonad.

### Neofunctionalization of ovary-promoting genes and new genetic pathways are implicated in ovary-testis transformation

Many paralogous genes, arising from an ancient teleost-specific whole genome duplication (*30*), show divergent sex-specific expression patterns during gonadal sex change (Fig. 3C). These include critical female-pathway genes, such as *foxl2* and genes in the Rspo1/Wnt/β-catenin signaling pathway known to regulate ovarian differentiation in mammals (*31, 32*). *Wnt4* (wingless-type MMTV integration site family, member 4), which activates *Ctnnb1* (β-catenin) and *Fst* (follistatin) to maintain mammalian ovarian development (*33*) (Fig. 3A), is duplicated in the bluehead wrasse: *wnt4a* was downregulated early along with *cyp19a1a,* consistent with a conserved feminizing role, while its paralogue *wnt4b* was sharply upregulated in late sex change and is expressed only in mature testes (Fig. 3C). Furthermore, *rspo1* (R-spondin-1), which stimulates *Wnt4* in mammals (Fig. 3A), also showed increasing expression during testicular construction, and multiple follistatin-like genes (e.g., *fstl4*) showed opposing sex-specific expression patterns (Fig. 3C). Neofunctionalization of duplicated sex-pathway genes may underpin the notable sexual plasticity of teleost fish.

Another unexpected result from our analysis was the progressive upregulation of genes involved in the JAK-STAT signaling pathway, from stage 2 to 5 (Table S2). JAK-STAT signaling plays an important role in determination and maintenance of male sexual fate in *Drosophila melanogaster* (*34, 35*), but has not previously been implicated in vertebrate sexual development.

### Cortisol pathways show dynamic expression across sex change

Cortisol, a glucocorticoid that controls stress-induced responses in all vertebrates, may be an important mediator of environmental sex determination (*36*). We therefore determined if cortisol-pathway genes show altered expression during early sex change stages. We find that the 11β-hydroxylase gene *cyp11c1,* responsible for cortisol production, is upregulated in bluehead wrasse gonads from stage 2. Furthermore, two genes encoding 11*β*-hydroxysteroid dehydrogenases (11*β*-HSD), *hsd11b2* and *hsd11b1-like,* show differential expression across sex change, but in opposite directions (Fig. 4A). The products of these genes mediate the stress response by respectively producing cortisol from inactive cortisone and *vice versa*, and their expression patterns imply a shift from cortisol production to inactivation from stage 3. We also observed opposite expression changes in genes encoding the glucocorticoid and mineralocorticoid receptors, *nr3c1* and *nr3c2* (nuclear receptor subfamily 3 group C members 1 and 2), respectively (37); *nr3c2* matched *hsd11b1la* expression changes, while *nr3c1* followed *hsd11b2* (Fig 4A). Together, high expression of *cyp11c1, hsd11b1la* and *nr3c2* at stage 2 (Fig. 4A) implies a window of high cortisol activity at the beginning of sex change, coincident with aromatase silencing and early ovarian atresia. Our data therefore suggest dynamic cortisol production and signaling at early sex change may be a key factor triggering female to male transition in the bluehead wrasse.

### Ovary-to-testis transformation involves extensive epigenetic reprogramming through a developmentally potent intermediate

Epigenetic marks such as DNA methylation and trimethylation of histone H3 lysine 27 *(H3K27me3)* are required during specification of pluripotent cells in mammalian embryos and the subsequent maintenance of cell identity in adulthood (*38, 39*). They are also increasingly recognized as key mediators of sexual differentiation in non-mammals (*19, 20, 40*). To explore their role in bluehead wrasse sex change, we investigated the expression profile of their regulatory machinery. We found that *ezh2, suz12, eed* and their co-factors *jarid2* and *rnf2,* components of the polycomb-repressor group 2 (PRC2) complex responsible for creating *H3K27me3* in vertebrates (*41, 42*), were downregulated during early sex-change and then reactivated following the transition to a masculine phenotype (Fig. 5C). *Kdm2aa,* a histone demethylase gene associated with transcriptional repression, had the opposite expression pattern, implying that, at least from an epigenetic perspective, the wrasse intermediate gonad takes on characteristics akin to the early developmental cells of mammals (*38, 39*). Furthermore, histone variants such as *h2az,* which feature in pluripotent stem cells, also showed their highest expression at stage 3, dropping immediately at stage 4 when testicular development starts.

**Fig. 5.**
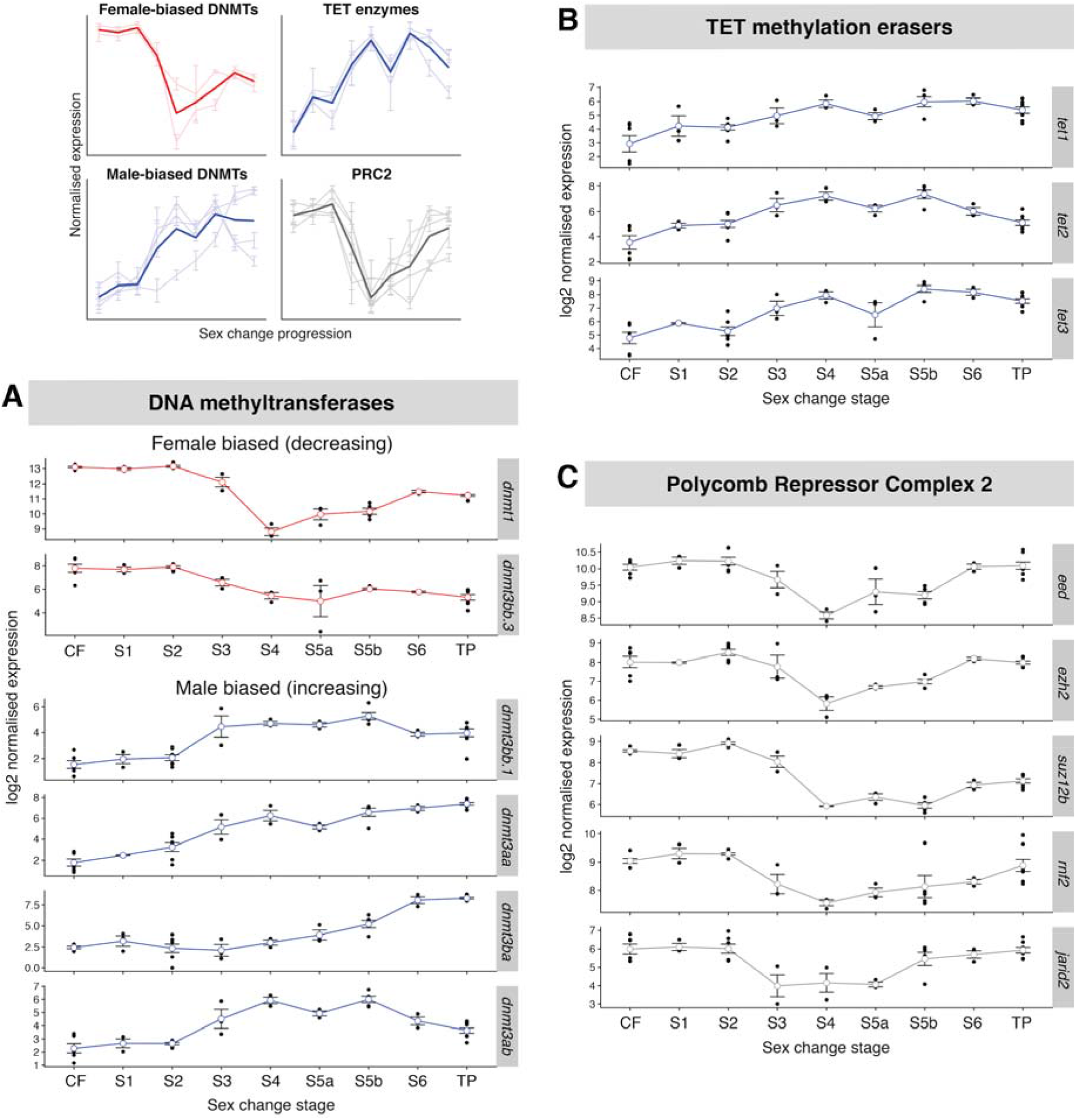
Epigenetic factors orchestrate sex change. **(A)** Normalized gonadal expression of DNA methyltransferase genes, showing a turn-over in sex-specific expression across sex change. The maintenance methyltransferase orthologue *dnmt1* is female-biased and has lowest expression at mid-sex change, whereas most methyltransferase *dnmt3* orthologues, responsible for *de novo* methylation, show increasing expression towards maleness. **(B)** Ten-eleven translocation (TET) methylcytosine dioxygenases, which demethylate 5-methylcytosines (5mCs), show highest expression during late sex change. **(C)** Genes encoding proteins *(eed, ezh2, suz12b)* and co-factors *(rnf2, jarid2)* of the chromatin remodeling Polycomb Repressor Complex 2 (PRC2) are suppressed during mid sex-change.

In mammals, global DNA demethylation features not only in naïve pluripotent stem cells, but also primordial germ cells (PGCs) – the bipotential germline progenitors of egg and sperm (*43*). Mammalian DNA demethylation in naïve stem cells and PGCs is driven by downregulation of *de novo* methyltransferase 3 (DNMT3) genes, deactivation of maintenance methyltransferase activity (DNMT1), and overexpression of the Ten Eleven Translocation (TET) family of DNA demethylation enzymes (*44–46*). We found that midway through sex change, there was a peak in TET expression (Fig. 5B). This indicates that like H3K27 modification, there is an intense period of DNA methylation reprogramming at the intermediate stages of sex change. Unexpectedly, we found that not only was there reduced *de novo* methylation machinery at intermediate stages of sex change, but actually wholesale replacement of sex-specific DNMT3 orthologues during the transition from ovary to testis (Fig. 5A). The *dnmt3bb.3* gene (orthologue of mammalian *Dnmt3b)* showed female-specific expression that declined during sex change and was replaced by male-specific expression of *dnmt3aa, dnmt3ab, dnmt3ba* and *dnmt3bb.1* (orthologues of mammalian *Dnmt3a* and *Dnmt3b).* While some sex-biased expression of DNMT genes has been observed in mammals (*47*), such distinct sex-specific expression appears to be unique to fish. Divergence in roles of *dnmt3* paralogues are described during development in zebrafish (*48, 49*), but this is the first report of sex-specific roles for DNA methyltransferase genes.

### Sex change involves genome-wide methylation reprogramming

In order to characterize the epigenetic effect of DNA methylation machinery replacement during sex change, we performed low-coverage bisulfite sequencing of DNA derived from the gonads of control females, TP males and transitioning individuals (Fig. 6). We found that ovary and testis showed significantly different CG methylation levels (70.7% and 82.1%, respectively), with gonads undergoing masculinization progressively accumulating methylation (Fig. 6A). To confirm this effect was specific to the gonad, we also tested brain tissue and found that CG methylation was relatively low (64.1%) and not sexually dimorphic.

**Fig. 6.**
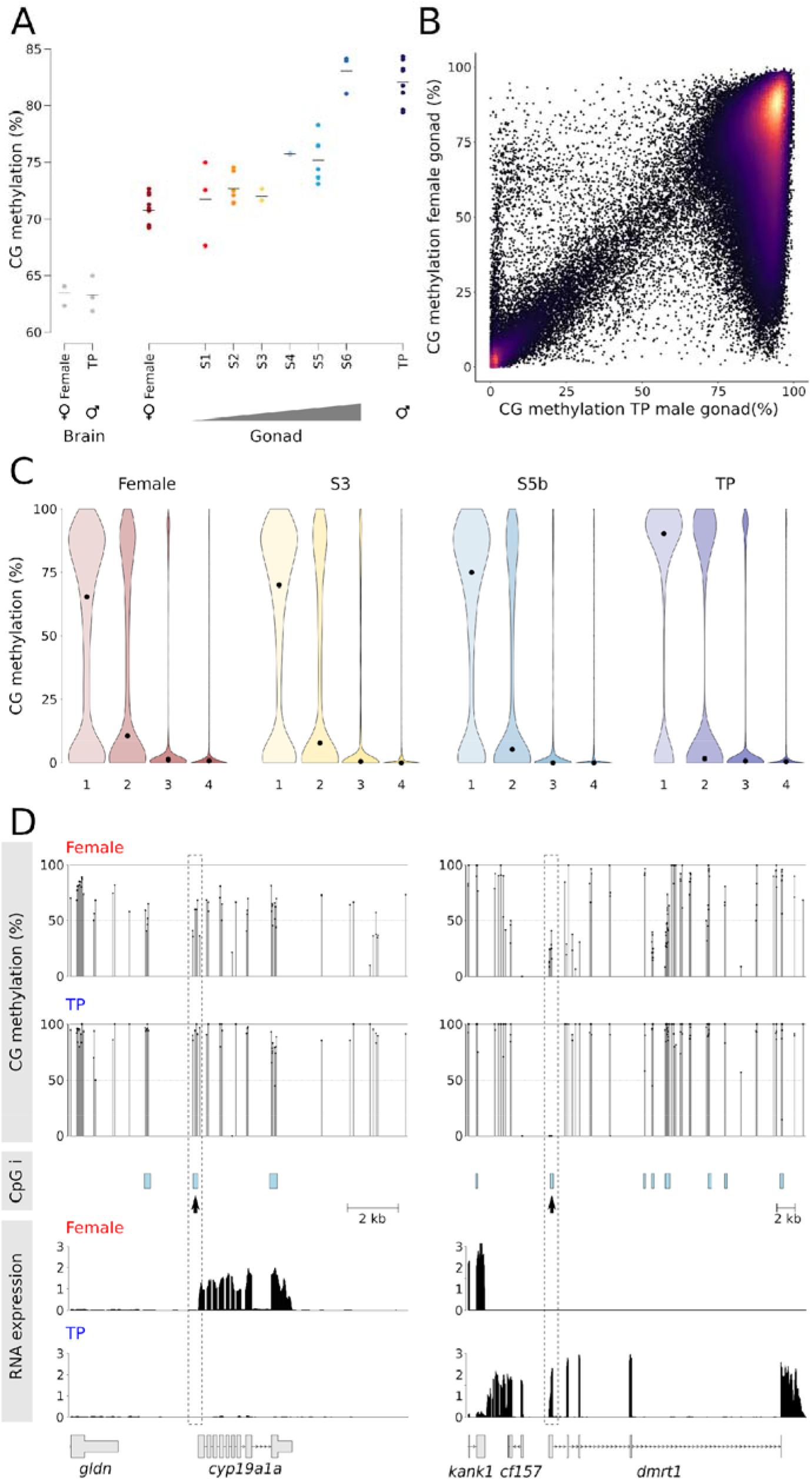
Global methylation changes and relationship between methylation and gene expression during sex change. **(A)** Global CG methylation levels during sex change examined by low-coverage sequencing. The horizontal bar indicates the mean; grey dots, brain samples; colored dots, gonadal samples. **(B)** Comparison of methylation (2 kb running windows) between female and terminal phase (TP) male gonadal methylomes. Only probes with > 100 calls were included for the analysis. **(C)** Violin plot showing distribution of methylation at transcription start sites of genes classified into quintiles according to expression level (4 = highest). Each violin is scaled to the same maximum width (total area is not constant between violins) to demonstrate distributions for each quintile. Black dots denote the median. **(D)** Relationship between CpG methylation and RNA expression for *cyp19a1a* and *dmrt1.* CG methylation track shows methylation levels for dinucleotides with > 10 calls, CpG islands (CpGi) predicted according to Gardiner-Garden and Formmer criteria. Changes in TSS methylation containing CpG islands (dashed box) are negatively correlated with gene expression for both genes (lower panel).

In order to characterize where global methylation changes were occurring during sex change, we undertook deep sequencing of selected female, intermediate and TP male libraries and compared this to a draft *de novo* assembly of the bluehead wrasse genome (25). When 2-kb running windows were analyzed genome wide, a striking number of low- and intermediately-methylated regions in females became fully methylated in TP males (Fig. 6B).

When targeted to gene promoters, particularly those with enriched levels of CG dinucleotides (known as CpG islands, or CGIs), DNA methylation is associated with gene silencing throughout all jawed vertebrates (*50*). To explore what effect the global increase in DNA methylation has upon gene expression in bluehead wrasse during sex change, we binned genes into quartiles according to their expression levels and asked what levels of DNA methylation existed in their promoter regions (Fig. 6C). We found that DNA methylation and gene silencing were coupled in a similar fashion throughout sex change, meaning that despite the major increase in global DNA methylation during transition, DNA methylation has the capacity to enforce gene silencing at all stages of reprogramming.

The methylation patterns of key genes involved in the sex change process provides evidence for the role of DNA methylation reprogramming in gonadal transformation. A CGI linked to the aromatase *(cyp19a1a)* transcriptional start site was hypermethylated as gene silencing progressed during sex change (Fig. 6D). Reciprocally, as the *dmrt1* gene became activated in transitioning fish, a promoter-linked CGI was progressively demethylated.

## Discussion

Transcriptomic and methylome analyses across sex change in the iconic bluehead wrasse have identified the triggers of socially-induced sex change and enactors of gonadal metamorphosis. Our results suggest that the environmental stimulus is exerted via stress, the subsequent steps involve repression of aromatase, and that distinctive epigenetic reprogramming is associated with re-engineering ovaries into testes. Significantly, this does not occur by direct transdifferentiation, but involves an intermediate state with altered epigenetic machinery expression that is reminiscent of mammalian naïve pluripotent stem cells and PGCs.

We observed early change in expression of the cortisol pathway that mediates stress. Removing the dominant TP male from a social group probably imposes severe social stress on large females, who must now compete for the reproductively privileged position of dominant male. Within minutes, transitioning females display aggression and courtship behaviors typical of dominant males, along with a rapid temporary color display (a bluish coloring of the head and darkening of pectoral fin tips) that is used to establish dominance.

The stress response in sex-changing females is expected to elicit rapid signaling changes of brain neurotransmitters, such as arginine vasotocin (AVT), gonadotropin-releasing hormone (GnRH), and norepinephrine (NE), which have been suggested to control behavioral sex change in social wrasses (*36, 51, 52*). We observed no significant early expression changes in genes encoding these neuropeptides or their precursors in the brain, but this may be because the sex-specific signal is subtle (*53, 54*) or highly localized (55). However, we did observe upregulation of isotocin *(it),* the homologue of mammalian oxytocin, at stage 1. In fish, *it* is linked to territoriality and aggression (*56*), is upregulated in response to cortisol (*57*), and so may facilitate early dominance establishment in transitioning females (Fig. 7). We also found that gonadal expression of dopamine beta-hydroxylase (dbh), which converts DA to NE (*58*), is up-regulated at stage 1 and downregulated at stage 2, consistent with the idea that stress-induced changes in NE signaling can trigger sex change in wrasses (*59*).

**Fig. 7.**
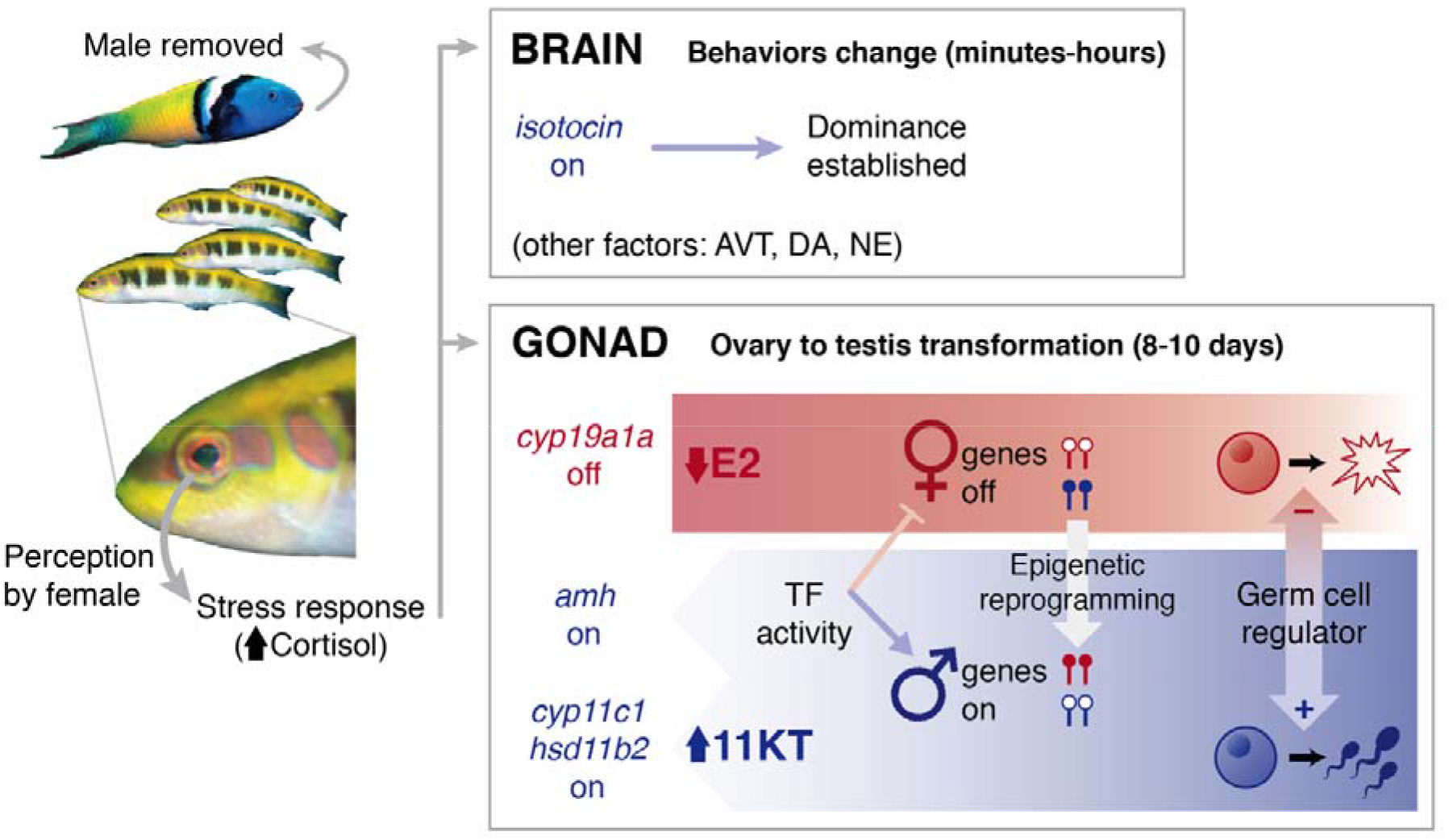
Mechanistic hypothesis of socially-cued sex change. Perception of a social cue (absence of a dominant male) promotes sex change in the largest female of a social group via raised cortisol. In the brain, cortisol increases *isotocin* expression to promote male-typical behaviors that rapidly establish social dominance. Other neuroendocrine factors may play a role (e.g., arginine vasotocin, AVT; dopamine, DA; norepinephrine, NE), although expression changes were not seen for their encoding genes. In the gonad, cortisol promotes transition from ovary (red) to testis (blue) via three pathways; 1) downregulates aromatase *(cyp19a1a)* expression causing estrogen (E2) production to cease and feminizing expression to decline, causing ovarian atresia, 2) upregulates *amh* expression, which as a transcription factor (TF) and germ cell regulator can suppress feminizing genes and promote oocyte apoptosis while promoting masculinizing expression and spermatogonial recruitment, and 3) upregulates androgenesis genes *cyp11c1* and *hasd11b2* to increase 11-Ketotestosterone (11-KT) production and support testicular development. Epigenetic reprogramming, via changes in sexually dimorphic DNA methylation (represented by lollipops, open = unmethylated, filled = methylated), re-writes cellular memory of sexual fate and associated sex-specific expression.

Cortisol can then initiate gonadal sex change, and our data implicate pathways that promote stress-induced masculinization of genetic females in artificial settings (Fig. 7): 1) suppression of aromatase expression via glucocorticoid response elements in the *cyp19a1a* promoter, 2) upregulation of *amh* expression to induce germ cell apoptosis and promote maleness, and 3) cross-talk with the androgenesis pathway via increased *cyp11c1* and *hsd11b2* expression (dual roles in 11-KT synthesis and cortisol metabolism) (*36, 60*). Our finding of opposing expression patterns for genes encoding glucocorticoid and mineralocorticoid receptors (*nr3c1* and *nr3c2*) and the enzymes that control cortisol production *(cyp11c1* and *hsd11b1la)* and inactivation *(hsd11b2),* implies highest cortisol production in early sex change (stage 2) (Fig. 4), consistent with observations in fishes and other vertebrates undergoing natural or temperature-induced sex reversal (*19, 61*). Therefore, in vertebrates where the environment exerts an influence on sex, cortisol might critically link environmental stimuli with sexual fate by initiating a shift in steroidogenesis.

Following the dramatic downregulation of aromatase and estrogen production, we observed a steady collapse of the female network before a male-promoting network was progressively established. Although previous work on protogynous wrasses (*62*) and protandrous clown fish (*63*) are equivocal about the importance of the FOXL2 and DMRT1 transcription factors in the sex change process, our data show they are important later, and are not triggers of sex change.

Neofunctionalization, where a gene homologue acquires novel function following gene duplication, is also readily evident in our data. Important sex-pathway genes are duplicated in the bluehead wrasse and appear to have acquired new, sometimes-unexpected roles. While duplicated copies of male-promoting genes (e.g., *sox9)* appear to have retained male-specific functions, many duplicated female-promoting genes have homologues showing a complete reversal in sex-specific expression (e.g., *foxl2, wnt4, fstl)* (Fig. 3C). In particular, key components of the Rspo1/Wnt/β-catenin signaling pathway, which regulates ovarian fate in mammals, are duplicated with one homologue showing upregulated expression during mid-to-late sex change when testicular structures are forming or have formed. This expression pattern suggests such duplicates have acquired new roles associated with male sexual fate, notably testicular differentiation. This flexibility in roles suggests a less conserved female genetic network operates in bluehead wrasses, and potentially other teleosts.

The use of neofunctionalized paralogues in sex change was not restricted to hormonal and signaling pathways; we also found duplicated epigenetic machinery in bluehead wrasse that exhibited female- or male-specific expression. Global DNA methylation was remodeled as expression of female-specific *de novo* methyltransferases was replaced with male-specific expression. The peak in TET expression seen at this time indicates remodeling of DNA methylation that is typical of mammalian PGCs and both naïve and classically grown pluripotent stem cells (*38*). The same appeared to be true for histone modifying machinery - the PRC2 complex is associated with differentiation in mammals, and showed high expression in cells belonging to committed sexual phenotypes but was deactivated in the gonads of transitional fish. Likewise, a histone demethylase and variant histones showed high expression in transitional fish. While alternative splicing of the histone methyltransferase *jarid2* has been associated with environmental sex-reversal in a dragon lizard (*19*), our study is the first to show comprehensive replacement of epigenetic machinery during sex change in a vertebrate, and that this is analogous to epigenetic reprogramming in germline and pluripotent cells of mammals. We hypothesize that sex-specific changes in the expression of epigenetic machinery is central to plasticity of sexual phenotype seen in sex-changing fishes.

Sexual fate has long been assumed to be canalized and stable throughout life, as it generally is for mammals and birds. However, manipulation of genes in the sex determining pathway of mammalian models now shows that gonadal sex requires active maintenance via antagonistic genetic signaling to suppress pathways of the opposite sex into adulthood (*12, 13, 64*). Thus, sexual fate may be inherently phenotypically plastic in all vertebrates, not just sex-changing fish (*65*).

In summary, this study reveals how environmental factors trigger gonadal sex change via a genetic cascade that re-engineers an ovary into a testis through an epigenetically distinct intermediate state. These findings enhance our understanding of how tissue reprogramming is controlled at the most fundamental level, and also shed light on the evolution of sex determination and differentiation mechanisms in vertebrates.

## Supporting information

## Acknowledgments

We thank Bill Tylor, Sidney Gaston Sanchez, Brandon Klapheke, Jeannie Brady, and Alison Lukowsky for support collecting samples, Jodi Thomas for assistance preparing DNA, Simon Fisher for aesthetic advice on Fig. 7, Mark Lokman, Dean Jerry and Francesc Piferrer for feedback on an earlier draft, and members of the Gemmell Lab for support.

## Author contributions

NJG, JRG, HL, EVT, TAH and OOR conceived the ideas and designed the study. JRG, MSL and HL collected the samples. HL, MSL, OOR and OK performed the laboratory work. OOR, EVT, HL and TAH performed the bioinformatic analyses, with support from KMR, HC and MAB. HC performed the genome scaffolding and annotation. JAM, TAH, EVT, HL and NJG wrote the manuscript and all authors contributed editorially and approved the final version;

## Data and materials availability

Sequencing data and the draft bluehead wrasse genome assembly are available from NCBI under BioProject PRJNA293777. Scripts used in the reported analyses are available on GitHub (https://github.com/hughcross/bluehead_methylome_bioinformatics/).

## Supplementary Materials

Materials and Methods

Figures S1-S2

Table S1

External Databases S1-S2

References (*66–104*)

