## Supplementary material for "Stress, novel sex genes and epigenetic reprogramming orchestrate socially-controlled sex change"

**This PDF file includes:**

Materials and Methods

Figs. S1 to S2

Table S1

Captions for Data S1 to S2

**Other Supplementary Materials for this manuscript include the following:**

Data S1. RNA-seq metadata for bluehead wrasse brain and gonad samples

Data S2. Whole-genome bisulfite sequencing of bluehead wrasse gonads

**Materials and Methods:**

**Experimental Design**

Female bluehead wrasses were induced to change sex by removing dominant terminal phase (TP) males from established social groups on patch reefs off the coast of Key Largo, FL, in May 2012 and May-June 2014 (*22, 66*). On each reef, all initial phase (IP) females and males larger than 45 mm standard length were captured and sexed by examination of the sexually dimorphic genital papilla and extrusion of gametes by gentle abdominal pressure. Females were tagged and returned to home reefs with TP males. IP males were relocated to distant reefs. Two days following tagging, TP males were removed to allow females to compete for dominance and undergo sex change. Females exhibiting TP-male-typical behaviors (*22*) were captured at increasing time points to produce a time-series of samples across the sex-change process. Tagged females showing no signs of behavioral sex change (as verified by histology) were captured as controls. All samples were collected around the daily spawning period.

Fish were euthanized with an overdose of MS-222 (Sigma) within two minutes of capture and the brain and gonads dissected immediately. The brain and one gonadal lobe were preserved in RNAlater (Life Technologies, Inc.) on ice, followed by storage at -20 ºC overnight, then -80 ºC until RNA extraction. The second gonadal lobe was fixed for histological analysis in 4% paraformaldehyde/1X PBS overnight at 4 ºC, followed by storage in 1X PBS before fixation in paraffin for histological sectioning with hematoxylin and eosin staining (Histology Laboratory, College of Veterinary Medicine, NCSU). Experiments were approved by the Institutional Animal Care and Use Committee at North Carolina State University (NCSU).

Gonadal sections were examined under light microscope to determine sex change status. In total, 41 samples were partitioned into successive stages (Fig. 1) based on gonadal histology (*23*) and behaviors observed at time of capture (*22*). Females showing no signs of behavioral or gonadal sex change (healthy ovaries with mature follicles and intact zona pellucida) served as control females. Behavioral and histological characteristics of sex change stages are summarized in Fig. 1. Sex changers at stage 5 (ongoing spermatogenesis) were further divided into stage 5a and stage 5b to reflect their divergent global gene expression patterns (Fig. 2B). Sample sizes for each stage were as follows: six control females (CF), three stage 1 (S1), seven stage 2 (S2), three stage 3 (S3), three stage 4 (S4), three stage 5a (S5a), five stage 5b (S5b), three stage 6 (S6) and eight TP males.

**RNA Sequencing and Whole Genome Bisulfite Sequencing (WGBS)**

Brain and gonadal tissues were homogenized using TissueLyser II (Qiagen) and total RNAs were extracted with TriReagent (Invitrogen), using chloroform (forebrain/midbrain) or bromochloropropane as the phase separation reagent. RNA samples were column-purified with either a NucleoSpin RNA XS kit (Macherey-Nagel) after DNase treatment (TURBO DNA-free Kit, Ambion) (2012 samples), or a Total RNA Purification Kit (Norgen Biotek) with on-column DNase digestion (RNase-free DNase I Kit, Norgen Biotek) (2014 samples). Only the forebrain/midbrain were used for RNA extraction (removing the hindbrain containing corpus cerebelli, pons, and medulla) as these contain regions belonging to the social behavior network and mesolimbic reward system, two neural circuits that are involved in the regulation of social-decision making (*55*) and likely to be key integrators and drivers of socially-induced sex change.

Total RNA concentration was measured by Qubit 2.0 Fluorometer (Life Technologies). RNA integrity was assessed on an Agilent 2100 Bioanalyzer. Sex-changing gonads consistently showed RNA profiles with a strong peak of low molecular weight RNA, which possibly corresponds to massive 5S rRNA expression in atretic ovaries and masks the 18S and 28S rRNA peaks used for calculating RNA integrity numbers (RIN) in teleost ovaries and intersex gonads (*67, 68*). Therefore, RIN values could not serve as useful measures of RNA integrity in sex-changing gonads of bluehead wrasses, although only samples with visibly intact rRNA peaks were used.

Library preparation and RNA sequencing were performed by the Otago Genomics and Bioinformatics Facility at the University of Otago under contract to New Zealand Genomics Limited. Samples were prepared as individual Illumina TruSeq Stranded mRNA libraries. The 2012 and 2014 samples were sequenced, with multiplexing, on separate occasions. For samples collected in 2012 (3 control females, 3 TP males, 6 intersex fish), 100 bp PE reads were generated over 4 lanes of an Illumina HiSeq 2000. For the 2014 samples (3 control females, 5 TP males and 21 intersex fish), 125 bp PE reads were generated over 1.5 lanes of an Illumina HiSeq 2500. Sequencing results including SRA accessions are summarized in Data S1.

Genomic DNA for WGBS was extracted from the TRIzol-homogenized samples used for RNA sequencing following manufacturer’s instructions, with post-extraction clean-up by magnetic beads (*69*). DNA was extracted from the brain and gonad of additional animals collected in Belize (2013) and Florida (2016) using a dual RNA/DNA Purification kit (Norgen) or a lithium chloride protocol (*70*).

WGBS was undertaken using a post-bisulfite adapter tagging (PBAT) method adapted from (*71*). Briefly, bisulfite treatment was performed with an EZ Methylation Direct Mag Prep kit (Zymo Research) following manufacturer’s instructions. Bisulfite treatment is performed prior to adapter tagging enabling simultaneous DNA fragmentation and conversion of unmethylated cytosines. Sequencing adapters are then added by complementary strand synthesis using random heptamer priming, and finally unique molecular barcodes and sequences necessary for binding to Illumina flow-cells are added to libraries by PCR.

Initially, we measured global DNA methylation by performing low-coverage sequencing of 6 brain (female and TP male) and 38 gonad samples (all stages) using a single-end MiSeq 100 bp protocol (Illumina) until the desired depth (at least 10,000 mapped CG calls) was attained (*50*). Subsequently, we performed deep sequencing on a subset of 17 gonad samples on a HiSeq 2500 instrument (Illumina) using rapid run mode, to obtain full methylomes and nucleotide-level methylation data for replicate females, TP males, and sex-changers (stage 2 to 5). Detailed sequencing and CpG quantification results are provided in Data S2.

**Statistical Analysis**

*Expression quantification*

Read quality was assessed in FastQC v0.11.5 (http://www.bioinformatics.babraham.ac.uk/projects/fastqc). Raw reads were trimmed of sequencing adaptors and low-quality bases (PHRED < 5) using Cutadapt v1.16 (*72*). Expression estimates for each sample were obtained using the ‘align_and_estimate_abundance.pl’ script within the Trinity package v2.6.6 (*73*): for each library, trimmed reads were mapped against our published bluehead wrasse transcriptome assembly (*24*) using Bowtie2 v2.3.2 (*74*) and transcript abundances were estimated using RSEM v1.3.0 (*75*) with default settings. Transcript-level count matrices were generated for brain and gonad samples separately using Trinity’s ‘abundance_estimates_to_matrix.pl’ script and the scaledTPM method.

*PCA and gene set enrichment*

Principal component analysis (PCA) (*76*) (centered, unscaled) was used to visualize transcriptome-wide expression variation within groups and among samples, following normalization by variance stabilizing transformation in DESeq2 v1.20.0 (*77*) in R v3.5.0 (*78*) 2018. The top 10,000 transcripts with greatest variance across samples were used and confidence ellipses around barycenters were plotted using the stat_conf_ellipse() function in ggpubr v0.1.8. For gonad only, to identify the 500 transcripts contributing most to each principal component (percentile 5th and 95th), component loadings (defined as eigenvectors scaled by the square root of the respective eigenvalues) were represented as coordinates in a Cartesian plane. Given the bimodal and skewed distribution of the values, percentiles rather than standard deviations were used as thresholds. Thresholds were defined as the 5th and 95th percentiles and divided the plane into four spatial regions: ‘Female’ and ‘Male’ represented the extremes of PC1, sexually ‘Differentiated’ and ‘Transitionary’ represented the extremes of PC2. Among regions, unique and shared transcripts were represented in a Euler diagram using the R package eulerr v4.1.0. To validate the results of our method, we visualized normalized expression across sex change for unique transcripts from each spatial region.

To identify the functional categories of genes uniquely associated with each spatial region, GO term enrichment analysis (*79*) was performed using TopGo v2.32.0 (*80*). and zebrafish genome annotations downloaded from Bioconductor (*81*) (org.Dr.eg.db v3.6.0). Significant GO terms (P < 0.01) for “Biological Processes” and “Molecular function” were identified using Fisher’s exact test with “weight01” algorithm. To reduce GO term redundancy and summarize the results, GO terms and P values weighted based on the scores of neighboring GO terms were used as input for REViGO (*82*), using *SimRel* as a semantic similarity measure, medium allowed similarity (0.7), and the *D*. *rerio* GO database.

*Differential expression and enrichment analyses*

Differential expression analyses were performed for brain and gonad separately using a generalized linear model (GLM) framework in DESeq2. Differentially expressed transcripts were called using the Wald test in pairwise comparisons between sex change stages and control females, and between neighboring stages, after fitting a single GML to estimate size factors and dispersion across all samples per tissue. False Discovery Rate (FDR) was controlled at 5% to account for multiple testing, and an adjusted significance value (FDR-P) of <0.05 was used to define significant differential expression. For gonadal samples, only transcripts with a log_2_ fold-change (LFC) >1 were considered. No fold change cut-off was applied in analyses of brain samples, due to the relatively subtle expression differences observed.

Transcripts showing differential expression at each sex-change stage (compared to control females) and in comparisons between neighboring stages were searched against Ensembl zebrafish protein database (BLASTX, E-value cut-off: 10-10) (*83*). Matched zebrafish protein IDs were converted to unique Ensembl zebrafish gene IDs via BioMart (*84*), and used for gene pathway over-representation analysis in DAVID v6.8 (*85, 86*).

*WGBS analysis and correlation with RNA-seq*

Raw WGBS sequences were processed in TrimGalore! v0.4.5 (https://www.bioinformatics.babraham.ac.uk/projects/trim_galore/); sequencing adapters were removed and 10 bp was trimmed from the 5’ end of reads to account for sequence biases associated with PBAT library construction, followed by removal of low-quality base calls (Phred score <20). Read mapping and base calling was performed in Bismark v0.19.0 (*87*) specifying the option --pbat. Our draft bluehead wrasse genome assembly (below) was used as a reference, obtaining an average of 51.47% mapping efficiency (sd. ± 3.07). BAM files were deduplicated and reports containing CG methylation were generated. The bisulfite treatment non-conversion rate was evaluated with the frequency of non-CG methylation, with all libraries having a conversion efficiency of at least 98.94% (Data S1).

CG methylation calls were analyzed in SeqMonk v1.42.0 (https://www.bioinformatics.babraham.ac.uk/projects/seqmonk/). Genome scaffolds were grouped into 23 pseudo-chromosomes and tracks were built using in-house annotations (details below). Among-sample variation was examined with PCA analysis, using 10 kb probes generated in non-overlapping windows with a minimum of 100 CG calls. To compare among stages, the PCA was re-run with individual samples merged and using 2 kb windows with minimum 100 CpG calls.

To examine methylation across transcription start sites (TSS), the longest transcript per gene was used. TSS were defined as 200 bp centered on the first nucleotide of an annotated mRNA, and a minimum of five methylation calls was applied as a threshold for inclusion. To evaluate coupling between TSS methylation and gene expression, trimmed RNA-seq reads were mapped to the reference genome using HTseq v0.9.1 (*88*) and imported into SeqMonk, specifying a minimum mapping quality of 60 to select only uniquely aligned reads. The SeqMonk RNA-seq quantitation pipeline was used to generate raw counts across exons of protein coding genes. Counts were corrected by transcript length and DNA contamination, and transcripts were divided into quintiles according to expression level.

To analyze methylation at individual CGs, probes of 2 consecutive nucleotides with a minimum of ten methylation calls were generated and the percentage methylation measured as number of methylated calls / total calls. CpG islands were identified using published methodology (*89*). For the regions of interest, 200 bp windows moving at 1 bp intervals were considered CpG islands if the Obs/Exp value was greater than 0.6 and the GC content exceeded 60%.

Scatter plots and violin plots were drawn using ggplot2 v3.0.0 in R (*90*). Histograms and genome annotations were generated using Gviz v1.24.0 (*91*).

*Bluehead wrasse draft genome assembly, scaffolding and annotation*

To provide a genomic reference for our methylation analyses, we constructed the first genome assembly for the bluehead wrasse. Ovary tissue of a single adult female (sex verified by histological analysis), from which methylome data was also derived, was used to provide DNA for sequencing. Genomic DNA was isolated using a lithium-chloride protocol (*70*), including RNAse digestion, followed by column purification through a DNA Clean & Concentrator kit (Zymo Research). Two TruSeq PCR-free libraries (350 and 550 bp inserts) and two Nextera mate-pair libraries (5 and 8 kb inserts) were constructed and sequenced together on 2 lanes of an Illumina Hi-Seq Rapid V2 (2x250 bp PE) at the Otago Genomics and Bioinformatics Facility at the University of Otago. Sequencing yielded a total of 1.84×10^11^ bases of data: 271.1, 182.4, 139.9 and 143.6 million paired reads from the 350 bp and 550 bp TruSeq, and 5 kb and 6 kb Nextera Mate Pair libraries, respectively. Based on the unassembled data, genome size was estimated at 0.76 Gb using SGA preqc (*92*).

Separate assemblies were performed for each TrueSeq library using the DISCOVAR de novo assembler (*93*) with default parameters. Based on the higher number of input reads and post-assembly statistics, the 350 bp insert assembly was used as a substrate for scaffolding. Prior to scaffolding, reads were trimmed of sequencing adapters and low-quality bases in Trimmomatic v0.35 (*94*), using the parameters ‘TRAILING:26’ and ‘MINLEN:20’.

Scaffolding was performed in SSPACE v3.0 (*95*) using trimmed data from all four sequencing libraries. First, reads from both TruSeq libraries were mapped to the DISCOVAR contigs in BWA (*96*). Mate Pair libraries were pre-processed with NextClip v1.3.1 (*97*) and all category A, B, and C mapping files were used for scaffolding. Resulting bam files were sorted by sequence name and converted to tab-delimited SSPACE files using the in-built script ‘sam_bam2tab.pl’. SSPACE was run several times for parameter optimization, with the following parameters used for final scaffolding: ‘minimum number of links to consider read pair (-k)’, 3; ‘maximum ratio between best two contig pairs (-a)’, 0.7; ‘minimum overlap between contigs to merge (-n)’, 15; and contig extension: ‘minimum number of reads needed to call a base during extension (-o)’, 10; ‘minimum number of overlapping bases during overhang consensus buildup (-m)’, 50; ‘minimal base ratio to accept overhang consensus base (-r)’, 0.8.

GapFiller v1-11 (*98*) was used to close gaps in scaffolds. Three iterations were run, using data from all four libraries and the following parameters: ‘minimum number of overlapping bases (-m)’, 50; ‘minimum number of reads to call a base (-o)’, 2; ‘minimal base ratio to accept overhang consensus (-r)’, 0.7; ‘minimum overlap to merge two sequences (-n)’, 10; ‘number of nucleotides trimmed at sequence edges of the gap (-t)’, 10.

Annotation was performed following the Trinotate version 3.1.1 (*99*) and PASA version 2.2 (*100*) pipelines. To first improve gene models from previous transcriptome annotations, and determine gene positions across the genome, transcripts from our published transcriptome assembly for bluehead wrasse (*24*) were aligned to the genome assembly using GMAP version 2018-03-11 (*101*) and BLAT version 3.5 (*102*), and these results were then fed into PASA. The program seqclean version x86_64 (https://sourceforge.net/projects/seqclean/files/) was used to validate the transcript sequences and trim unwanted sequences (e.g., vectors, adaptors, polyA tails). Using both the cleaned and original transcripts, several rounds of PASA were performed, incorporating the genome alignments, and our previous transcriptome annotations (*24*).

The Trinotate pipeline was used to annotate the mapped assemblies from the PASA output. First, TransDecoder version 5.0.2 (https://github.com/TransDecoder) was used to predict coding regions. Predicted peptide sequences and transcripts were then used as queries to search multiple protein and nucleotide databases. The protein database SwissProt was queried using BLAST (*103*), using *blastp* for peptide sequences and *blastx* for transcripts. Additionally, protein databases of the Zebrafish and Tilapia genomes were searched. The program HMMER 3.1b2 (http://hmmer.org/) was used to identify protein domains in the peptide sequences using the Pfam database. Search results were consolidated into a Trinotate sqlite database to produce an annotation report.

Custom Python and R scripts were used to extract annotations from the Trinotate report and link these to the mapped location of transcripts on the genome (<https://github.com/hughcross/bluehead_methylome_bioinformatics>). A custom Python script was also used to create separate mapping files (gff3 format) for the three gene references used (sprot, zebrafish, tilapia), using the best match annotation as the gene description. Mapping files were used to visualize gene annotations in SeqMonk.

The scaffolded assembly was more complete and more contiguous than the DISCOVAR assembly and was used in methylation analyses. The scaffolded genome included 379,332 scaffolds, with a scaffold and contig N50 of 15.6 ad 12.5 kb, respectively, and total length of 1095.9 Mb. According to a BUSCO analysis (*104*), this assembly is relatively complete (96.7% complete, 1.4% fragmented orthologues), but with a duplication level of 13.2%. Although the large number of scaffolds indicates a fragmented assembly, over 91% of the total genome length is represented within large scaffolds (29,971 over 1 kb in length; 10,270 over 10kb in length). Therefore, although scaffolds less than 1 kb in length were numerous (349,361; 92.1%) these represent less than 9% of the genome length and, from initial surveys, largely represent repetitive regions. Furthermore, almost no genes mapped to these small scaffolds. Therefore, only scaffolds larger than 1 kb were used for bisulfite mapping and methylation analyses.

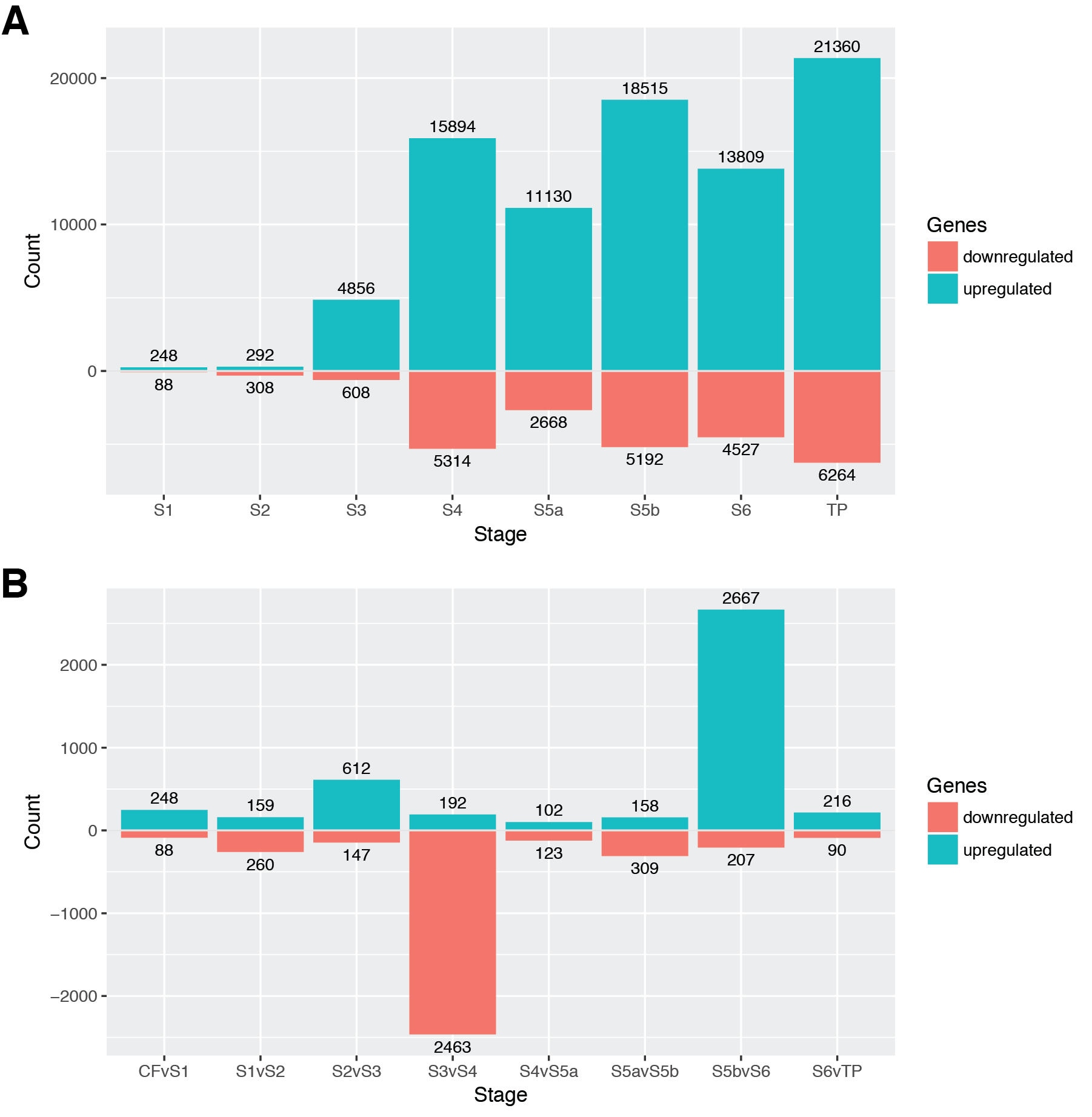

**Fig. S1.** Differential transcript expression in gonad of bluehead wrasses across sex change. Numbers of differentially expressed transcripts in pairwise comparisons between (**A**) control females and sex-change stages and (**B**) neighboring stages. Cut-off: adjusted p-value <0.05 and fold-change >2. Up/down-regulation refers to the second stage in each comparison. CF, control female; S, stage; TP, Terminal Phase male.

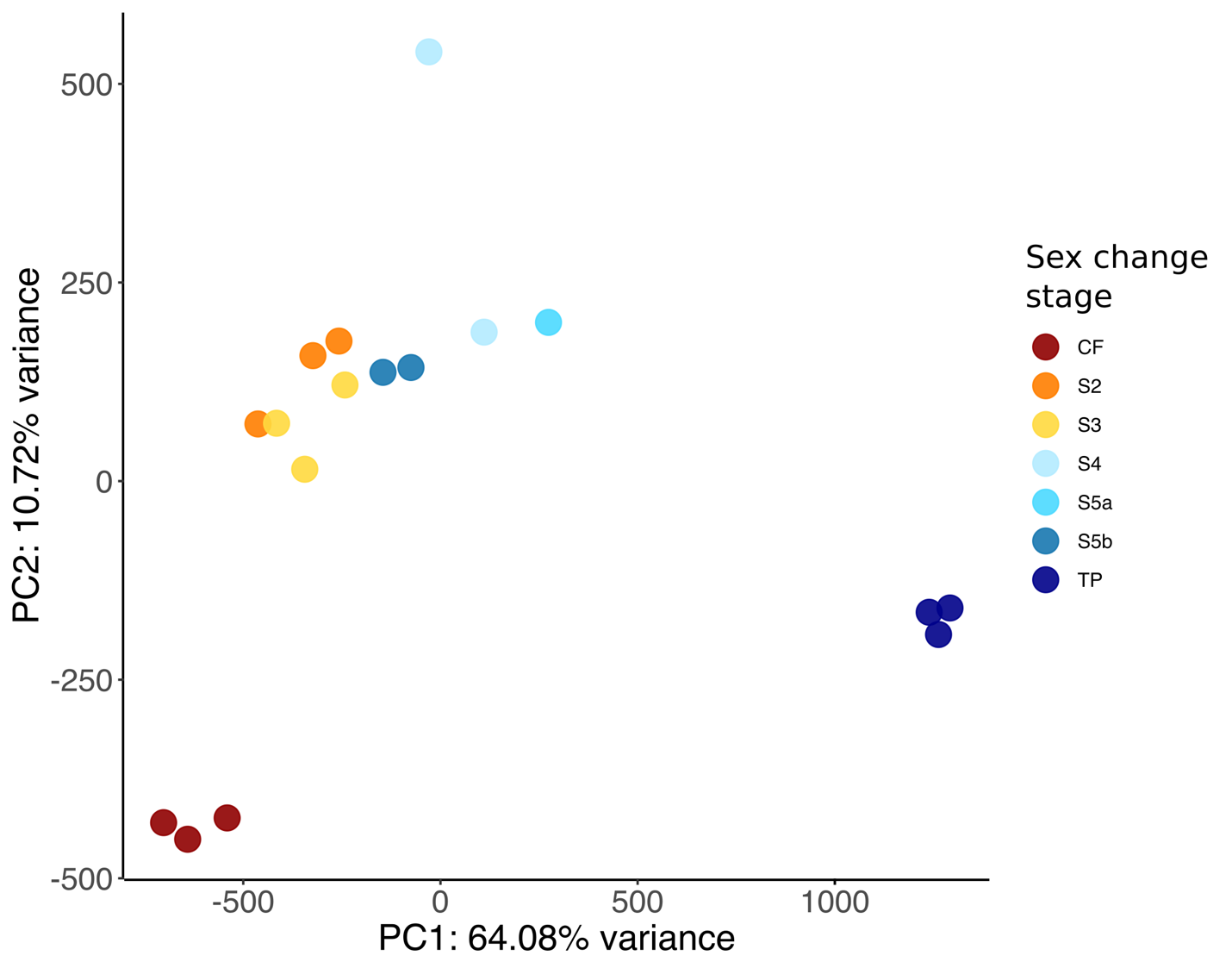

**Fig. S2.** Principal component analysis (PCA) of gonadal methylomes during sex change in bluehead wrasse. For the analysis only probes of 10 kb with more than 100 CpG calls were included. CF, control female; S, stage; TP, Terminal Phase male.

| **Comparison** | **CF v S2** | **CF v S3*** | **CF v S4*** | **CF v S5a*** | **CF v S5b*** | **CF v S6** |
| --- | --- | --- | --- | --- | --- | --- |
| **Number of genes** | 2 | 20 | 41 | 32 | 37 | 27 |
| **Fold enrichment** | na | 1.9 | 1.7 | 1.7 | 1.5 | 1.4 |
| **P-value** | na | 7.60E-03 | 1.90E-04 | 2.20E-03 | 7.80E-03 | 8.10E-02 |
| **Upregulated genes enriched in   Jak-STAT signaling pathway** |  |  | *akt3b* | *akt3b* | *akt3b* | *akt3b* |
|  |  |  |  |  |  | *bcl2l1* |
|  |  | *ccnd2a* | *ccnd2a* | *ccnd2a* | *ccnd2a* | *ccnd2a* |
|  | *cish* | *cish* | *cish* | *cish* | *cish* |  |
|  |  | *cntfr* | *cntfr* | *cntfr* | *cntfr* |  |
|  |  |  | *crfb4* | *crfb4* | *crfb4* | *crfb4* |
|  |  | *csf2rb* | *csf2rb* | *csf2rb* | *csf2rb* | *csf2rb* |
|  |  | *csf3r* | *csf3r* | *csf3r* | *csf3r* | *csf3r* |
|  |  | *epor* | *epor* | *epor* | *epor* | *epor* |
|  |  |  | *ghra* | *ghra* | *ghra* |  |
|  |  |  | *grb2b* |  |  |  |
|  |  |  | *ifngr1l* |  | *ifngr1l* |  |
|  |  |  | *il10* |  | *il10* |  |
|  |  | *il11ra* | *il11ra* | *il11ra* | *il11ra* | *il11ra* |
|  |  |  | *il13ra1* |  | *il13ra1* |  |
|  |  | *il13ra2* | *il13ra2* | *il13ra2* | *il13ra2* | *il13ra2* |
|  |  | *il21r.1* | *il21r.1* | *il21r.1* | *il21r.1* | *il21r.1* |
|  |  |  | *il22ra2* |  |  |  |
|  |  | *il2rga* | *il2rga* | *il2rga* | *il2rga* | *il2rga* |
|  |  |  | *il6* | *il6* |  |  |
|  |  | *il6st* | *il6st* | *il6st* | *il6st* |  |
|  |  |  | *il7r* | *il7r* | *il7r* |  |
|  |  |  | *irf9* | *irf9* | *irf9* |  |
|  |  | *jak1* | *jak1* | *jak1* | *jak1* | *jak1* |
|  |  |  | *jak2b* |  | *jak2b* |  |
|  |  | *lifra* | *lifra* | *lifra* | *lifra* | *lifra* |
|  |  | *m17* | *m17* | *m17* | *m17* | *m17* |
|  |  |  |  |  |  | *pik3ca* |
|  |  |  | *pik3cd* | *pik3cd* | *pik3cd* | *pik3cd* |
|  |  |  | *pik3cg* |  |  | *pik3cg* |
|  |  |  | *pik3r1* | *pik3r1* | *pik3r1* |  |
|  |  |  | *pik3r2* | *pik3r2* | *pik3r2* | *pik3r2* |
|  |  | *pik3r5* |  |  |  |  |
|  |  |  | *pik3r3b* | *pik3r3b* | *pik3r3b* | *pik3r3b* |
|  |  |  | *pik3r5* | *pik3r5* | *pik3r5* | *pik3r5* |
|  |  |  | *pim1* | *pim1* | *pim1* |  |
|  |  |  |  |  |  | *pim2* |
|  |  |  |  |  | *ptpn11b* | *ptpn11b* |
|  |  | *ptpn6* | *ptpn6* | *ptpn6* | *ptpn6* | *ptpn6* |
|  |  | *si:dkey-13m1.2* | *si:dkey-13m1.2* | *si:dkey-13m1.2* | *si:dkey-13m1.2* | *si:dkey-13m1.2* |
|  |  | *socs1a* | *socs1a* | *socs1a* | *socs1a* | *socs1a* |
|  |  | *socs3b* | *socs3b* | *socs3b* | *socs3b* |  |
|  |  |  | *socs7* |  | *socs7* |  |
|  |  |  | *stam2* |  |  | *stam2* |
|  | *stat1b* | *stat1b* | *stat1b* | *stat1b* | *stat1b* | *stat1b* |
|  |  |  | *stat4* | *stat4* | *stat4* |  |

**Table S1.** Genes upregulated in sex-changers (stage S2 to S6) against control females (CF) and enriched in the Jak-STAT signaling pathway. *Indicates significant enrichment (P-value <0.05) in a DAVID functional enrichment analysis.

**Data S1. RNA-seq metadata for bluehead wrasse brain and gonad samples (separate file)**

The table lists the number of trimmed reads, and average quality ‘Q’ value obtained for each RNA-seq library and used in downstream analyses, and the corresponding NCBI BioSample and SRA accession number/s. *RIN values for ovarian samples are not representative of RNA quality as the large quantity of small RNAs interferes with calculations. N/A not calculated. ^Number of read pairs after quality trimming and filtering, used in expression analyses.

**Data S2. Whole-genome bisulfite sequencing of bluehead wrasse gonads (separate file)**

The table lists the number of cytosine calls at either symmetric CG dinucleotides ('CG') or in other sequence contexts ('non-CG'), mapped against the draft bluehead wrasse genome. Number of calls are following deduplication. The frequency of non-CG methylation indicates the maximum rate of non-conversion during the bisulfite treatment step; by this measure, all libraries had a bisulfite conversion efficiency of at least 98.94%. na, not applicable.
